# SpoT and GppA hydrolases prevent the gratuitous activation of RelA by pppGpp in *Escherichia coli*

**DOI:** 10.1101/350843

**Authors:** Rajeshree Sanyal, Rajendran Harinarayanan

## Abstract

Stringent response, a conserved regulation seen in bacteria, is effected through the modified nucleotides (p)ppGpp. The metabolic cycle of these molecules is driven by the synthase activity of RelA and SpoT and the hydrolase activity of SpoT and GppA which together sets the basal (p)ppGpp pool. Growth arrest due to (p)ppGpp accumulation from basal RelA activity apparently explained the essentiality of SpoT hydrolase function. We found, pppGpp degradation was enhanced when the SpoT hydrolase activity was lowered or eliminated and when this was alleviated by inactivation of the GppA hydrolase, gratuitous synthesis of (p)ppGpp by RelA was activated, leading to growth arrest. The RelA-ribosome interaction was not mandatory for these phenotypes. Our results show, for the first time, elevated pppGpp promoted the amplification of RelA-mediated stringent response in the absence of established RelA activating signals in the cell and the SpoT and GppA hydrolases prevented this. The accumulation of pppGpp inhibited the SpoT hydrolase activity. We propose this autocatalytic activation of RelA by pppGpp is likely to be an allosteric regulation and can result in a bistable switch.

## Introduction

Intracellular signaling molecules play a key role in the physiology of organisms by regulating key cellular processes and coordinating them with extracellular or intracellular signals (Pesavento and Hengge, 2009). In eubacteria, (p)ppGpp is a signaling molecule that accumulates during starvation, switching the balance of metabolism from growth and cell division to survival and stress response (Chatterji and Kumar Ojha, 2001; Braeken *et al.*, 2006; Potrykus and Cashel, 2008; Hauryliuk *et al.*, 2015). (p)ppGpp, also sometimes referred to as an ‘alarmone’, consist of a pair of molecules - guanosine tetraphosphate (ppGpp) and guanosine pentaphosphate (pppGpp), which are synthesized by the transfer of a pyrophosphate moiety from ATP to GDP or GTP respectively (Potrykus and Cashel, 2008; Atkinson *et al.*, 2011). The gram-negative bacteria *Escherichia coli* is a well-studied model organism with regard to the role of (p)ppGpp in bacterial physiology. β-and α-proteobacteria, including *E. coli*, have two proteins which are involved in stress response and (p)ppGpp metabolism - RelA and SpoT (Mittenhuber, 2001; Atkinson *et al.*, 2011). Both proteins are members of the Rel/Spo homolog (RSH) family and share similar domain architecture. The N-terminal half of the protein has the catalytic domain with the (p)ppGpp synthase and hydrolase functions in the case of SpoT and only synthase function in case of RelA. The C-terminal half of the protein has the regulatory domains important for sensing stress and starvation signals. RelA is a ribosome-bound protein that is activated by the “hungry” codons that appear following amino acid starvation and the consequent increase in the cellular concentration of uncharged t-RNA (Wendrich *et al.*, 2002). Recent cryo-electron microscopy studies have provided important insights into the structural basis for RelA activation by the entry of uncharged tRNA into the A-site of an elongating ribosome (Arenz *et al.*, 2016; Brown *et al.*, 2016; Loveland *et al.*, 2016). SpoT has a weak synthetic activity and a strong (p)ppGpp hydrolase activity and is an essential gene (An *et al.*, 1979; Xiao *et al.*, 1991). It not only regulates the basal (p)ppGpp levels in the cell (Sarubbi *et al.*, 1988), but also responds to various stress and starvation signals such as carbon (Xiao *et al.*, 1991), fatty acid (Seyfzadeh *et al.*, 1993) and iron (Vinella *et al.*, 2005) limitation. The SpoT hydrolase activity was inhibited in the presence of uncharged tRNA and more severely in the presence of ribosomes (Richter, 1980), conditions that mimic amino acid starvation. Various factors interact with SpoT and regulate the balance between its synthase and hydrolase functions. CgtA, a G-protein interacts with SpoT to up-regulate its hydrolase activity under nutrient-rich condition (Jiang *et al.*, 2007). The acyl carrier protein was reported to interact with SpoT and up-regulates its synthase activity during fatty acid starvation (Battesti and Bouveret, 2006). In addition to SpoT, pppGpp is hydrolyzed by GppA, a pentaphosphate phosphohydrolase that converts it into ppGpp (Somerville and Ahmed, 1979; Harat and Sy, 1983; Keasling *et al.*, 1993). The physiological relevance of this reaction is not clear. ppGpp being the predominant stringent nucleotide in the cell is considered to be the principal effector of stringent response (Mechold *et al.*, 2013).

The primary and well-studied target for (p)ppGpp is RNA polymerase, to which it binds independently or aided by the protein factor DksA (Ross *et al.*, 2013; Zuo *et al.*, 2013; Ross *et al.*, 2016) and alters the global transcriptome so as to promote survival under starvation or stress (Durfee *et al.*, 2008; Traxler *et al.*, 2012). Various models exist to explain the effects of (p)ppGpp at the level of transcription initiation or sigma factor competition (Barker *et al.*, 2001; Paul *et al.*, 2004; Magnusson *et al.*, 2005). Together with DksA, (p)ppGpp inhibits transcription of rRNA operons and ribosomal protein genes and activates amino acid biosynthetic genes (Paul *et al.*, 2004; Paul *et al.*, 2005). Studies have shown alternative targets for (p)ppGpp in *E. coli* and in other organisms (Kanjee *et al.*, 2012). This includes enzymes from the nucleotide biosynthesis pathway, GTPases, particularly ObgE, EFG, EFTu, RF3 and IF2 (Miller *et al.*, 1973; Hamel and Cashel, 1974; Rojas *et al.*, 1984; Milon *et al.*, 2006). DNA replication and cell division are regulated by (p)ppGpp (Schreiber *et al.*, 1995; Joseleau-Petit *et al.*, 1999; Wang *et al.*, 2007; Ferullo and Lovett, 2008).

Due to the essential nature of the SpoT function, studies have generally been performed using point mutants that altered the steady-state levels of ppGpp. Using a SpoT depletion system, changes in the (p)ppGpp metabolic pattern and the associated growth response can be monitored. In this study, we report, during SpoT depletion or in the hydrolase mutants of *spoT*, there was reduction in the pppGpp pool and this was required to support growth. Our results show, GppA and an uncharacterized activity together lowered the pppGpp pool and prevented the gratuitous activation of RelA-dependent (p)ppGpp synthesis that conferred growth arrest. Two hypomorphic *relA* alleles that allowed GppA-dependent survival in the absence of SpoT function were isolated and characterized. Using one of the alleles, we show that the accumulation of only ppGpp was insufficient to confer sustained growth arrest. Our results show pppGpp can serve as a positive signal for the amplification of RelA-mediated stringent response in the absence of known documented signals and therefore its level is tightly regulated by the SpoT and GppA hydrolases.

## Results

### Depletion of SpoT results in the enhanced degradation of pppGpp

The *spoT* gene function that is essential in the wild-type *E. coli* was dispensable in the *relA* mutant (Xiao *et al.*, 1991), and therefore, the essential *spoT* function was inferred as the degradation of (p)ppGpp synthesized through the basal RelA activity. To test this, (p)ppGpp accumulation was followed as the cellular SpoT activity was reduced using a system designed to gradually deplete it. To do this, the chromosomal *spoT* gene was knocked out and SpoT was expressed in a regulated fashion from a plasmid (see methods). For ease of genetic manipulations, either of the two null mutant alleles reported in the literature, namely, *spoT212*, a markerless *spoT* deletion or *spoT207::Cm* have been used in strain constructions. In the Δ*spoT*/pRCspoT strain wherein the chromosomal *spoT* gene was deleted, growth was expected to be dependent on the IPTG-driven expression of the *spoT* gene present in the single copy unstable plasmid *pRCspoT* (Nazir and Harinarayanan, 2016). When the Δ*spoT*/pRC*spoT* strain, cultured in the presence of IPTG was washed and sub-cultured, IPTG-dependent growth was observed (Fig. 1A) indicating that the system can be used to study the consequences of SpoT depletion.

**Figure 1.**
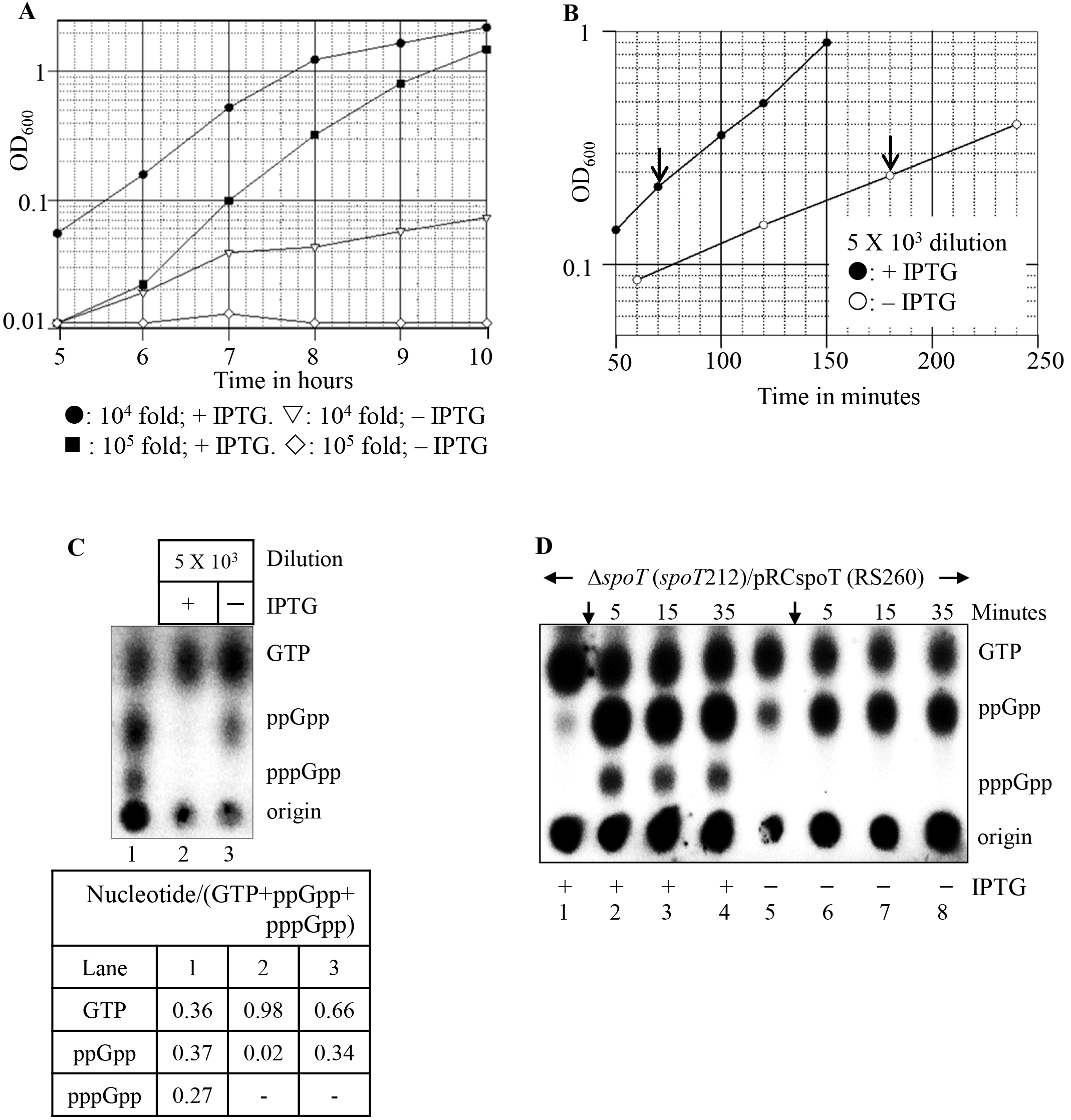
Depletion of SpoT is associated with growth retardation and accumulation of ppGpp, but not pppGpp. **A.** Δ*spoT*/pRC*spoT* (RS260) strain cultured in MOPS buffered medium with glucose, 20 amino acids, ampicillin and 1mM IPTG was washed and sub-cultured in the same medium with (•, ▪) or without (▿, ◊) IPTG at the dilutions indicated and the growth was monitored. **B.** Δ*spoT*::Cm/pRC*spoT* (RS14) was sub-cultured in the same medium used for Fig. 1A at the indicated dilutions. Growth after the subculture into the medium containing the ^32^P-H_3_PO_4_ is shown, the solid arrows indicate the time points at which the samples were collected for (p)ppGpp estimation. **C.** Samples collected as shown in Fig. 1B were subjected to PEI-TLC. Valine-induced stringent response in the wild type is provided as a reference (lane 1). Ratios of a nucleotide to the total (GTP+ppGpp+pppGpp) was calculated for each lane. **D.** Strain Δ*spoT*/pRC*spoT* (RS260) was cultured after 10^3^-fold dilution in MOPS glucose medium containing ampicillin and with (+) or without (−) IPTG, the stringent response was induced with valine (arrow), samples were collected immediately before valine addition or subsequently at the time indicated.

To monitor (p)ppGpp levels, cells cultured in the presence or absence of IPTG were allowed to undergo at least two divisions in the low phosphate medium containing ^32^P-H_3_PO_4_ and subjected to thin layer chromatography. This method allowed (p)ppGpp levels to be measured when the growth of the strain was retarded due to the depletion of SpoT (Fig. 1B and 1C). Following SpoT depletion, ppGpp accumulated, but pppGpp was not detectable. We tested different dilutions in order to deplete SpoT to varying extents, and in each case, the result was similar (data not shown). Unlike the Δ*spoT*/pRC*spoT* strain, the growth of the Δ*relA* Δ*spoT*/pRC*spoT* strain was less significantly inhibited following SpoT depletion (Fig. S1) and (p)ppGpp accumulation was not detected here (data not shown). Nevertheless, the growth rate of the strain was reduced in the absence of IPTG, which could be due to the composition of the growth medium used (all amino acids at 40μg/ml). It has been reported, the growth of the ppGpp^0^ strain in a defined medium was sensitive to amino acid composition (Potrykus *et al.*, 2011).

Due to the presence of amino acids in the growth medium, the signal for RelA activation would be low, this, and the conversion of pppGpp to ppGpp by GppA can together account for the lack of detectable pppGpp accumulation. To test this, isoleucine starvation was provoked by the addition of valine (Leavitt and Umbarger, 1961) before or after SpoT depletion and the (p)ppGpp synthesis was studied. In the culture induced for *spoT* expression, a typical stringent response, that is, accumulation of ppGpp and pppGpp and depletion of GTP was observed (Fig. 1D, lanes 1 - 4). This indicated, the SpoT expression from plasmid did not significantly perturb the stringent response. Following SpoT depletion, a significant accumulation of ppGpp was seen prior to starvation and this further increased with the onset of amino acid starvation; interestingly, pppGpp was still not detectable (Fig. 1D, lanes 5 - 8). This indicated, pppGpp accumulation was especially reduced, as compared to ppGpp during SpoT depletion. The absence of pppGpp could be due to its reduced synthesis or increased degradation or both. The results described below show the GppA hydrolase function was partly required to observe this phenotype. In early studies (Laffler and Gallant, 1974; Fiil *et al.*, 1977), it was reported that the hydrolase deficient *spoT1* allele accumulated mainly ppGpp during amino acid starvation. We confirmed that following amino acid starvation, the *spoT1* mutant, like the SpoT depleted strain, showed RelA-dependent accumulation of ppGpp, but not pppGpp (Fig. S2). These results suggest, counterintuitively, lowering the SpoT hydrolase activity lowered the pppGpp accumulation.

### Hypomorphic relA alleles provide evidence for concomitant synthesis and degradation of ppGpp during stringent response following amino acid starvation

To identify mutations that suppressed the growth defect of the Δ*spoT* strain, transposon-mediated mutagenesis was carried out in the Δ*spoT*::Cm/pRC*spoT* strain and mutants that survived the loss of pRC*spoT* were identified as white colonies from the LB IPTG X-Gal plates. In order to further identify the loss of function *relA* mutants amongst these, the white colonies were screened using the SMG plate test for RelA functionality (Uzan and Danchin, 1976). The *relA* null mutant or mutants that exhibit very low activity (such as *relA1*) do not grow on minimal medium with glucose and the amino acids serine, methionine, and glycine (SMG) (Fig. S3, rows 1 and 2). About 50% of the white colonies screened showed SMG sensitivity and were excluded from further analysis. The insertions in two separate SMG-resistant mutants were mapped and used for further studies. Somewhat surprisingly, one of the transposon insertions was within the *relA* gene (after the 496^th^ codon) and the other at the end of the *rlmD* gene that is immediately upstream of *relA* (Fig. 2A). These insertions, referred to as *relA*::Tn*10*dTet and *rlmD*::Tn*10*dKan, when individually introduced into the Δ*spoT*::Cm/pRC*spoT* strain by phage P1 transduction, allowed the segregation of white colonies on LB agar plates with IPTG and X-Gal (Fig. 2B, panels ii and iii), suggesting the presence of each mutation was sufficient to support growth in the absence of SpoT function. The *relA*::Tn*10*dTet Δ*spoT*::Cm and *rlmD*::Tn*10*dKan Δ*spoT*::Cm strains grew on minimal glucose medium with or without SMG, while the *spoT*^+^ derivatives did not grow in the presence of SMG (Fig. S3, rows 3, 4, 9 and 10). These results showed that a certain threshold of RelA activity and possibly (p)ppGpp accumulation can be tolerated in the absence of SpoT function. The *relA1* allele also survives the *spoT* deletion and the *relA1* Δ*spoT* strain grows in minimal medium (Xiao *et al.*, 1991) suggesting the presence of (p)ppGpp in this strain. *relA1* is a naturally selected hypomorphic allele present in the laboratory strains, wherein two polypeptides come together to reconstitute a weak (p)ppGpp synthase activity, which is evident upon over-expression (Metzger *et al.*, 1989). Unlike *relA1*, the *relA1* Δ*spoT* strain could grow slowly on SMG plate, with single colonies observed after 72 hrs (Fig. S4A). These results indicated that the basal (p)ppGpp pool increased in the absence of SpoT function in all the hypomorphic *relA* alleles tested.

**Figure 2.**
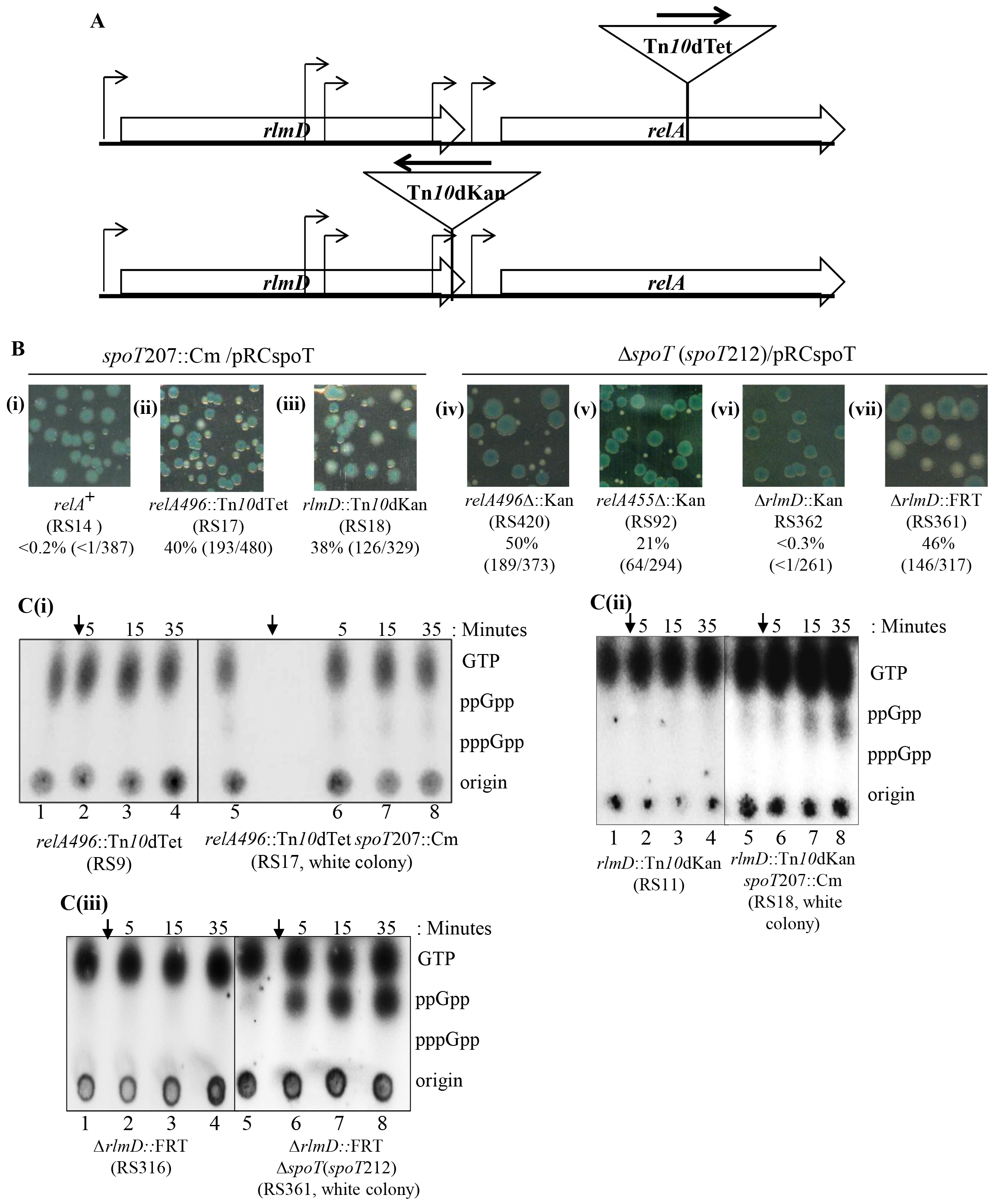
Hypomorphic *relA* alleles suppress the Δ*spoT* growth defect. **A.** Cartoon of the transposon insertions *rlmD*::Tn*10*dKan and *relA496*::Tn*10*dTet. Bent arrows represent known or annotated promoters, arrow within the transposon indicate the direction of transcription of the antibiotic marker. **B.** Segregation of the unstable plasmid pRC*spoT* was assayed as described in the methods. Relevant genotype of the strain, the percentage of white colonies and the total number of colonies (blue + white) used to calculate the ratio are indicated. **C.** Strains whose relevant genotypes are indicated were cultured in MOPS glucose medium, isoleucine starvation was induced with valine (arrow) and samples were collected immediately before valine addition and subsequently at the times indicated above the lanes and subjected to PEI-TLC. The line between lanes in the TLC indicates that internal lanes from a single TLC have been removed.

The transposon insertion after the 496^th^ codon of *relA* generated a UAA stop codon at the site of insertion and therefore was expected to produce a truncated polypeptide containing the 496 amino-terminal amino acids (RelA496Δ). A plasmid-encoded RelA polypeptide with the 455 amino-terminal amino acids, expressed from the IPTG-inducible P_tac_ promoter showed constitutive (p)ppGpp synthesis and was incapable of sensing amino acid starvation; the polypeptide had a half-life of 7.5 minutes as compared to the 2-3 hours for the full-length RelA protein (Schreiber *et al.*, 1991; Svitils *et al.*, 1993). To compare the phenotypes of the RelA truncations, precise deletions were engineered in the chromosomal *relA* gene. These alleles referred as *relA496*Δ::Kan and *relA455*Δ::Kan, when introduced in the Δ*spoT*/pRC*spoT* strain could support growth in the absence of SpoT function, which was evident from the appearance of white colonies (Fig. 2B, panel iv, and v) in the plasmid segregation assay. The white colonies with the *relA496*Δ::Kan Δ*spoT* and *relA455*Δ::Kan Δ*spoT* genotypes exhibited SMG-resistance unlike their *spoT*^+^ counterparts (Fig. S3, rows 5 - 8), and suggested that the absence of SpoT function elevated the cellular (p)ppGpp pool in these strains. Indeed, a slightly elevated basal ppGpp pool was observed in the SMG-resistant *relA*::Tn*10*dTet Δ*spoT*::Cm strain when compared to the SMG-sensitive *relA*::Tn*10*dTet *spoT*^+^ strain (Fig. 2C (i), lanes 1 and 5). Amino acid starvation did not elicit stringent response in either strain (Fig. 2C (i), lanes 2 - 4 and 6 - 8), and this is consistent with the reports that the RelA-CTD is required for sensing the ribosome-dependent amino acid starvation signals (Arenz *et al.*, 2016; Brown *et al.*, 2016; Loveland *et al.*, 2016). These results suggest, RelA’ (truncated RelA), when expressed at physiological levels could not elevate the (p)ppGpp pool in the presence of SpoT hydrolase activity.

The transposon insertion *rlmD*::Tn*10*dKan, 13-bp upstream from the *rlmD* stop codon (Fig. 2A) can potentially alter the RelA expression and also the RlmD activity. To test if the suppression of Δ*spoT* growth defect resulted from the loss of RlmD function, the Δ*rlmD*::Kan allele from the Keio collection (Baba *et al.*, 2006) was introduced in the wild-type and Δ*spoT*/pRC*spoT* strains by Phage P1 transduction. Suppression was not evident in the Δ*rlmD*::Kan Δ*spoT*/pRCspoT strain as it did not segregate white colonies on LB IPTG X-Gal plate (Fig. 2B, panel vi), however, flip-out of the Kanamycin marker supported the segregation of white colonies (Fig. 2B, panel vii) and the growth of these colonies on minimal medium containing SMG (Fig. S3, rows 11 - 13). These results showed that the loss of RlmD function was not necessary for suppression, while the altered *relA* expression after the flip-out of the Kan cassette was sufficient for the suppression of the Δ*spoT* growth defect. We propose the *rlmD*::Tn*10*dKan and Δ*rlmD*::FRT mutations reduced the *relA* expression, the former possibly terminated transcripts originating from the promoters within and upstream of *rlmD* (Metzger *et al.*, 1988; Nakagawa *et al.*, 2006; Brown *et al.*, 2014), and the latter eliminated the promoters reported within the *rlmD* ORF. Making use of *lacZ* reporter fusions at the chromosomal locus of these genes, the relative contribution of the promoters located within the *rlmD* ORF to those upstream was studied (Fig. S5). The promoters within the *rlmD* ORF increased the expression 75-fold, indicating that they predominantly contributed to the RelA expression. The reason, the Δ*rlmD*::Kan allele failed to suppress the Δ*spoT* growth defect could be because promoter(s) within the Kan^R^ cassette partially or fully compensated for the loss of expression from the native *relA* promoters.

SMG resistance was observed in the *rlmD*::Tn*10*dKan Δ*spoT*::Cm strain but not in the *rlmD*::Tn*10*dKan strain (Fig. S3, rows 9 and 10), however, no significant difference in basal ppGpp is evident between the two strains (Fig. 2C (ii), lanes 1 and 5). Similarly, the Δ*rlmD*::FRT strain showed SMG-resistance (Fig. S3, row 12), but increase in the basal ppGpp pool was not evident (Fig. 2C (iii) lane 1). Our results suggest, marginal increase in the ppGpp pool less than that which can be reliably detected by 1D-TLC was sufficient to confer SMG-resistance. The Δ*rlmD*::FRT Δ*spoT* strain showed SMG-resistance (Fig. S3, row 13) and a slightly elevated basal ppGpp pool (Fig. 2C (iii), lane 5). Amino acid starvation did not cause (p)ppGpp accumulation in the *rlmD*::Tn*10*dKan, *rlmD*::Tn*10*dKan Δ*spoT*::Cm and Δ*rlmD*::FRT strains (Fig. 2C (ii) lanes 2 - 4 & 6 - 8; Fig. 2C (iii) lanes 2 - 4). However, stringent response was elicited in the Δ*rlmD*::FRT Δ*spoT* strain and accumulation of ppGpp, but not pppGpp was observed (Fig. 2C (iii), 6 - 8); the rate of ppGpp accumulation was slower as compared to the wild-type or *spoT1* strain (compare Fig. 1D, lane 2; Fig. S2 lane 2 and Fig. 2C (iii) lane 6). This result provides evidence for the SpoT-mediated degradation of ppGpp being concomitant with the RelA-mediated synthesis of (p)ppGpp during stringent response. This was not entirely expected, because, the SpoT hydrolase activity was reported to be inhibited during the amino acid-induced stringent response (Richter, 1980). The concomitant synthesis and degradation of (p)ppGpp imply the concentration of these molecules at any given time is set by the sum total of their synthesis and degradation rates. This would have implications for the amplification of the stringent response, because, it was reported that (p)ppGpp increased the rate of their own ribosome-dependent synthesis by RelA *in vitro* (Shyp *et al.*, 2012; Kudrin *et al.*, 2018). The results imply, the SpoT hydrolase activity can directly contribute to the regeneration of GDP from stringent nucleotides during the stringent response and indirectly to GTP as it was proposed that GDP would be converted to GTP by Ndk (Kari *et al.*, 1977; Heinemeyer and Richter, 1978).

### GppA activity prevented gratuitous (p)ppGpp synthesis and growth arrest in strains with reduced spoT hydrolase activity

The accumulation of ppGpp but not pppGpp, after SpoT depletion or reduction in the hydrolase activity (*spoT1* mutant), suggested altered (p)ppGpp metabolism. One possibility was, RelA preferentially synthesized ppGpp under these conditions. Results in the previous section highlighted the role of SpoT hydrolase activity in the regeneration of GTP during the stringent response, therefore it was possible, pppGpp synthesis was limited by GTP availability. Furthermore, there was evidence *in vitro* that the *E. coli* RelA prefers GDP as a substrate over GTP (Sajish *et al.*, 2009). We, therefore, asked if increasing the cellular GTP pool in the *spoT1* mutant would support pppGpp accumulation during stringent response. We used the *gsk3* allele that codes for the feedback-resistant guanosine kinase and reported to increase the intracellular GTP pool in the presence of guanosine (Petersen, 1999). Normally there is a growth inhibition associated with GTP accumulation, but this can be overcome by histidine and tryptophan supplementation (Petersen, 1999). Stringent response in the *spoT1 gsk3* strain, in the absence of guanosine, was similar to that seen in the *spoT1* mutant. In the presence of guanosine, the only significant difference observed was the absence of GTP depletion (Fig. S6). Assuming this reflected an increase in the GTP pool, the result suggested an increase in the GTP level was insufficient to support pppGpp synthesis in the *spoT1* mutant.

Another possible reason for the absence of pppGpp accumulation could be, the pppGpp synthesized was rapidly turned over into ppGpp by GppA, a cytoplasmic pppGppase that is not ribosome-associated (Somerville and Ahmed, 1979). To test this, the Δ*gppA*::Kan allele, sourced from the Keio collection (Baba *et al.*, 2006), was introduced by phage P1 transduction to construct the Δ*spoT*::Cm Δ*gppA*::Kan/pRC*spoT* and *spoT1* Δ*gppA*::Kan/pRC*spoT* strains. During SpoT depletion, growth of the Δ*spoT*::Cm Δ*gppA*::Kan/pRC*spoT* strain was more severely inhibited as compared to the Δ*spoT*::Cm/pRC*spoT* strain (Fig. 3A (i)). The plating efficiency dropped at least three orders of magnitude in the former strain as compared to the latter strain (Fig. 3A (ii)). When the stringent nucleotides pool was determined in the Δ*spoT*::Cm Δ*gppA*::Kan/pRC*spoT* strain after the growth retardation that followed SpoT depletion (Fig. S7), accumulation of ppGpp, pppGpp, and depletion of GTP was observed (Fig. 3B). This showed the absence of pppGpp accumulation during SpoT depletion in the Δ*spoT*::Cm/pRC*spoT* strain was partly due to the GppA activity. In addition to pppGpp accumulation there was also a significant decrease in the GTP pool when compared to the SpoT depletion in *gppA*^+^ strain (compare Fig. 3B, lane2 with Fig. 1C, lane 3).

**Figure 3.**
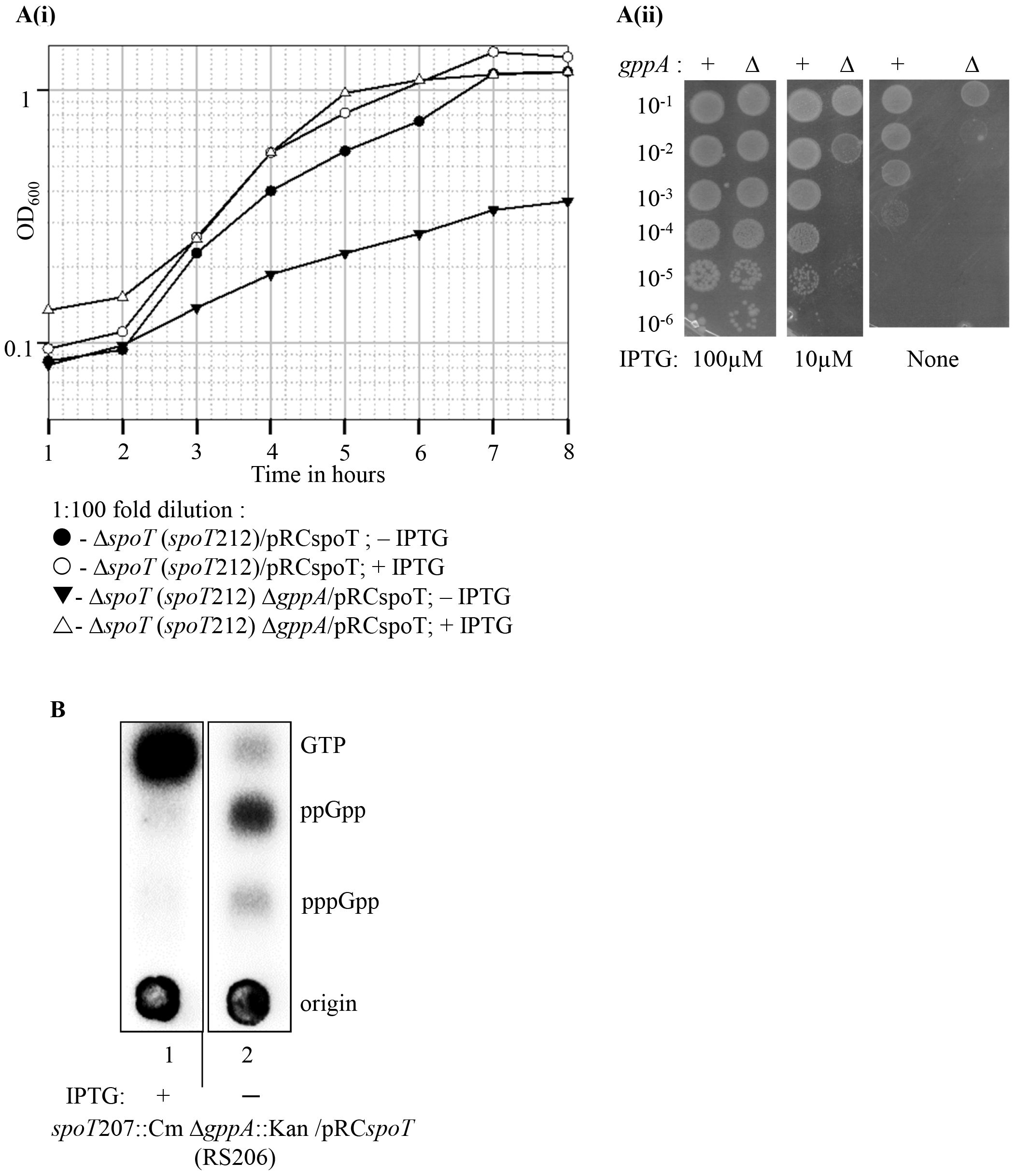
SpoT depletion in the *gppA* mutant was associated with severe growth inhibition, depletion of GTP and accumulation of ppGpp and pppGpp. **A. (i)**. Δ*spoT*/ pRC*spoT* (RS260) and Δ*spoT* Δ*gppA*::Kan/pRC*spoT* (RS478) strains cultured in the MOPS buffered medium containing glucose, 20 amino acids, ampicillin and 1mM IPTG were washed and sub-cultured in the same medium with (○,▵) or without (•,▾) IPTG at the dilutions indicated and the growth monitored **A. (ii)** Δ*spoT*::Cm/*pRCspoT* (RS14) and Δ*spoT*::Cm Δ*gppA*::Kan/*pRCspoT* (RS206) strains cultured in LB containing ampicillin and 1mM IPTG were washed, serially diluted in LB medium and the dilutions spotted on LB agar plates containing Amp and 100 μM, 10 μM or no IPTG and incubated for 16 hr at 37°C. **B.** Δ*spoT*::Cm Δ*gppA*::Kan/pRC*spoT* (RS206) cultured in the medium described in 3A (i) was diluted 200-fold in the same medium or without IPTG and sampled after at least two divisions in the presence of P^32^ (arrows in Fig. S7) and subjected to TLC. The samples were part of a single TLC with the lanes in between deleted.

We asked if the lack of pppGpp accumulation during the stringent response in the hydrolase deficient *spoT1* strain (Fig. S2) was also due to the GppA activity. Interestingly, synthetic growth defect was evident when the Δ*gppA*::Kan allele was introduced into the *spoT1* genetic background. The *spoT1* Δ*gppA*::Kan/pRC*spoT* strain failed to segregate white colonies (*spoT1* Δ*gppA*::Kan genotype) in the minimal glucose medium with or without casamino acids and the white colonies were slow growing in LB medium (Fig. 4A, panels ii, vi and x). Deletion of the *relA* gene suppressed the synthetic growth defect (white colonies in panels iv, viii and xii). The slow-growing *spoT1* Δ*gppA*::Kan strain could be maintained on LB plate but failed to grow on minimal glucose medium with or without casamino acids (data not shown). Apparently, an elevated basal pppGpp pool and lowered *spoT* hydrolase activity together conferred synthetic growth inhibition in the presence of RelA activity, while individually they affected the growth marginally at best (Fig. 4A, the white colonies in panels i, v and ix have the *spoT1* genotype, blue colonies in panels ii, vi and x have the Δ*gppA*::Kan genotype). Another hydrolase deficient *spoT* allele, *spoT202* (Sarubbi *et al.*, 1989), also showed synthetic growth defect with the Δ*gppA*::Kan allele and the growth defect was more severe than seen for the *spoT1* allele as no white colonies could be recovered on LB medium in addition to minimal medium (data not shown). This suggested, the synthetic growth defect with Δ*gppA* arose specifically from the SpoT hydrolase deficiency. As described below, the synthetic growth inhibition was associated with accumulation of (p)ppGpp and the reduced degradation of (p)ppGpp.

**Figure 4.**
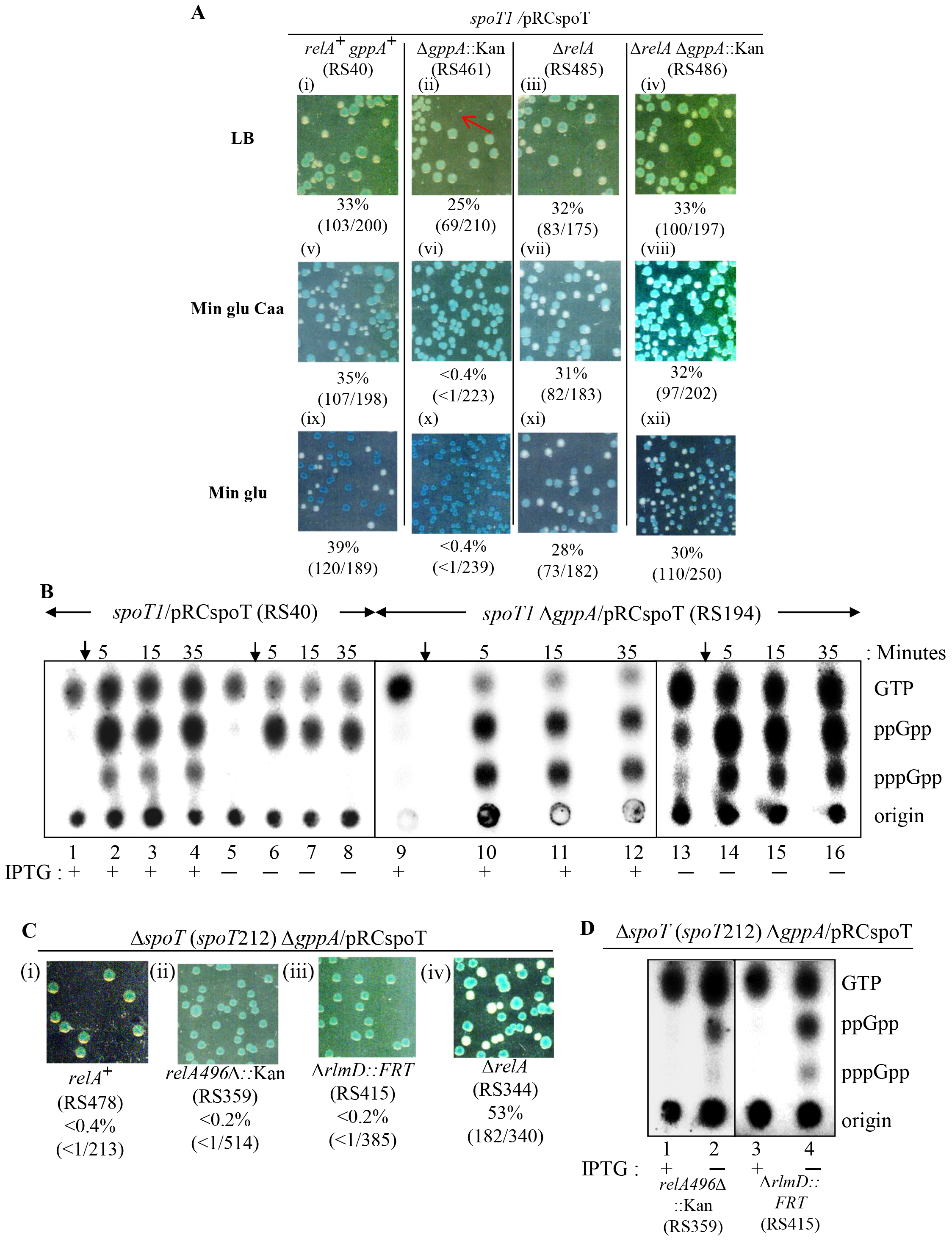
Lowering the SpoT hydrolase activity in the absence of GppA function conferred growth arrest from gratuitous RelA activation. **A.** The loss or retention of the unstable plasmid pRC*spoT* was assayed as described in the methods. Relevant genotype of the strain, the percentage of white colonies and the total number of colonies (blue+white) used to calculate the ratio are indicated. The panels within a column represent single genotype. LB and minimal glucose cas-amino acids plates were incubated at 37°C for 24 hours and minimal glucose plates for 48 hours. **B.** Stringent nucleotides pool was measured by TLC following growth in the presence of P^32^. Overnight cultures of *spoT1*/pRC*spoT* (RS40) (Lanes 1 - 8) and *spoT1* Δ*gppA*/ pRC*spoT* (RS194) (Lanes 9 - 16) were grown in MOPS medium containing glucose and ampicillin in the presence or absence of IPTG as indicated after 1in 1000 dilution. Amino acid starvation was induced with valine (arrow) and samples collected just before or subsequently at the indicated time. **C.** Plasmid segregation was studied in strains whose relevant genotypes are indicated and as described in methods. **D.** Stringent nucleotides pool was measured by TLC for the strains whose relevant genotypes are indicated. Cultures were inoculated at 1:1000 dilution in MOPS medium containing glucose, 20 amino acids and ampicillin with or without IPTG as indicated below the lanes and at least two doubling were allowed in the presence of P^32^.

To find out if the *spoT1* Δ*gppA*::Kan growth defect was accompanied by a change in the stringent nucleotides pool, SpoT depletion was carried out by IPTG withdrawal in the *spoT1* Δ*gppA*::Kan/pRC*spoT* strain. As a control, depletion was carried out in the *spoT1*/pRC*spoT* strain (Fig. 4B). Before and after SpoT depletion, the *spoT1*/pRC*spoT* strain exhibited stringent response like the wild-type and *spoT1* strains respectively. That is, isoleucine starvation induced ppGpp and pppGpp synthesis following growth in the presence of IPTG (Fig. 4B, lanes 1-4) and mainly ppGpp, following growth in the absence of IPTG (Fig. 4B, Lanes 5-8) similar to the *spoT1* mutant (Fig. S2, lanes 1 - 4). This indicated, IPTG withdrawal sufficiently depleted the plasmid-encoded protein that the phenotype of the chromosomal allele was evident. As expected, IPTG withdrawal also induced growth inhibition in the *spoT1* Δ*gppA*::Kan/pRC*spoT* strain, and associated with the growth inhibition the (p)ppGpp pools were elevated (Fig. 4B, lane 13) and which increased further following amino acid starvation (Fig. 4B, lanes 14-16). The results indicated that elevated pppGpp pool activates RelA-dependent (p)ppGpp synthesis in the *spoT1* strain without amino acid starvation. During stringent response in the presence of IPTG, the *spoT1* Δ*gppA*/pRC*spoT* strain accumulated similar levels of ppGpp and pppGpp (Fig. 4B, lanes 10 - 12) as reported for a *gppA* mutant (Somerville and Ahmed, 1979). However, after IPTG withdrawal, that is, in the phenotypically *spoT1* Δ*gppA* cells, pppGpp accumulated less than ppGpp (Fig. 4B, lanes 14-16), the plausible reasons for this are discussed later.

Since *relA* deletion suppressed the *spoT1 gppA* synthetic growth defect (Fig. 4A, panels iv, viii and xii), we tested the effect of the hypomorphic *relA* alleles. Plasmid-free derivatives of *relA496*Δ::Kan*spoT1* Δ*gppA*::Kan/pRC*spoT* and Δ*rlmD*::FRT *spoT1* Δ*gppA*::Kan/pRC*spoT* strains (white colonies) were recovered on LB and as well as minimal glucose medium with or without casamino acids (data not shown). This suggested wild-type level expression of full-length RelA was needed for gratuitous (p)ppGpp synthesis and growth arrest in the *spoT1* Δ*gppA*::Kan background. Since the *relA496*Δ::Kan and Δ*rlmD*::FRT alleles supported growth in the complete absence of *spoT* function (Fig. 2B, panels iv and vii) we asked if the growth of these strains were dependent on the GppA function. Plasmid free derivatives (white colonies) could not be recovered from the *relA496*Δ::Kan Δ*spoT* Δ*gppA*::Kan/pRC*spoT* and Δ*rlmD*::FRT Δ*spoT* Δ*gppA*::Kan/pRC*spoT* strains on LB (Fig. 4C) and in the minimal glucose medium as well (data not shown). SpoT depletion was carried out in these strains to ask if the growth arrest was accompanied by (p)ppGpp accumulation. Gratuitous synthesis of ppGpp and to a lesser extent, pppGpp, was observed in both strains (Fig. 4D). Notably, (p)ppGpp synthesis by the **relA496*Δ::Kan* allele indicated that ribosome binding was not necessary for the pppGpp dependent activation. These results indicate, the residual hydrolase activity in the *spoT1* allele (as compared to the Δ*spoT* allele) prevented the activation of hypomorphic *relA* alleles by pppGpp to gratuitously synthesize (p)ppGpp. We tested, if the *relA1* allele, possibly the weakest of the hypomorphic allele studied here also exhibited synthetic growth defect it the Δ*spoT* Δ*gppA*::Kan background. Plasmid-free derivatives could be recovered from the *relA1* Δ*spoT* Δ*gppA*::Kan/pRC*spoT* strain and no growth phenotype was evident in this strain as compared to the parental strain, *relA1* Δ*spoT*, in LB or minimal medium (Fig. S4B). This suggested, the *relA1* allele, unlike the *relA496*Δ or Δ*rlmD*::FRT alleles was not activated by pppGpp to carry out gratuitous (p)ppGpp synthesis. It is possible, in RelA1, the structural determinants required for allosteric activation by pppGpp may not be present since the (p)ppGpp synthase activity was reconstituted by two polypeptides in trans.

### SpoT mediated degradation of (p)ppGpp is inhibited by pppGpp

To understand the Δ*gppA spoT1* synthetic growth defect, the contribution of GppA and SpoT, individually and together, to the turnover of the stringent nucleotides was studied. Isoleucine starvation was induced with valine to allow the accumulation of (p)ppGpp and the kinetics of degradation was monitored after the reversal of starvation by isoleucine addition. As reported, the GTP pool decreased and that of the stringent nucleotides increased following amino acid starvation in the wild-type strain (Fig. 5A). In the Δ*gppA* strain, following starvation, the GTP and pppGpp pools were, respectively, 2.5-fold lower and 3.8-fold higher than in the wild-type; the ppGpp pool was not significantly altered (Fig. 5A & B, table S2). Consistent with the biochemical activity of GppA, an increase in the pppGpp pool was expected, but the decrease in GTP pool was unexpected. We suspected this could be due to the reduced regeneration of GTP from the stringent nucleotides following the inhibition of SpoT hydrolase activity by the increase in pppGpp. As seen from Fig. 2C (iii), SpoT activity contributes to the degradation of ppGpp to GDP, during amino acid starvation; GTP can then be made from GDP. Indeed, in the Δ*gppA* strain, following the inhibition of RelA activity by the reversal of amino acid starvation, we observed the rate of degradation of the stringent nucleotides and the accumulation of GTP was slightly reduced as compared to the wild-type strain (Fig. 5A & B, table S2). It is conceivable, pppGpp may be poorly hydrolyzed relative to ppGpp, by SpoT, and therefore, an increase in the pppGpp pool inhibited GTP generation by competitively inhibiting the ppGpp degradation to GDP by SpoT (Fig. 7).

**Figure 5.**
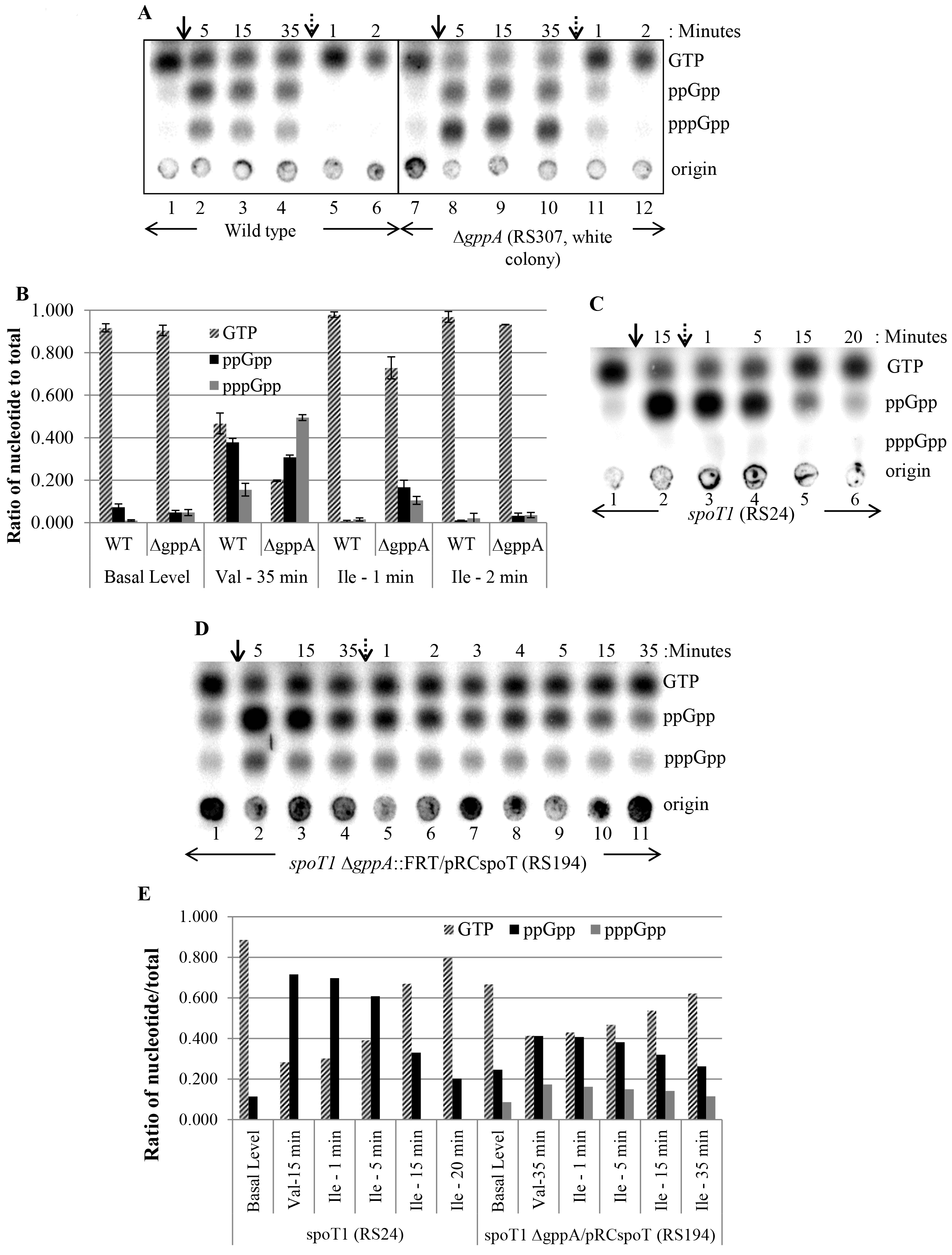
(p)ppGpp degradation is reduced by elevated pppGpp. **A.** TLC was performed for wild-type and Δ*gppA*::FRT (RS307, white colony) strains grown in MOPS glucose. Isoleucine starvation was induced with valine (arrow, solid line), and reversed by isoleucine addition (arrow, dotted line) and samples were collected at the indicated time and just before valine addition (lanes 1&7). **B.** The ratio of a nucleotide to total (GTP + ppGpp + pppGpp) was plotted for the samples collected before or after valine (Val) or after valine and isoleucine (Ile) addition. Time after valine and subsequent isoleucine addition are indicated below the bars. Data provided are the mean of two independent experiments along with the standard deviation of the mean. **C.** TLC was performed for the *spoT1* strain (RS24); growth condition and sample collection are as described in 5A. **D.** *spoT1* Δ*gppA*/pRC*spoT* (RS194) strain was grown in the MOPS glucose medium containing ampicillin and 1mM IPTG and sub-cultured 1 in1000 in the same medium but without IPTG. Following growth inhibition, TLC was performed as described in Fig. 5A. **E.** The ratio of a nucleotide to the total was plotted from the TLC’s shown in Fig. 5C and Fig. 5D. Sample collection times are indicated below the bars.

The effect of the increased pppGpp pool on (p)ppGpp degradation by the *spoT1* allele was studied. We confirmed the ppGpp degradation rate following amino acid starvation was reduced in the *spoT1* strain as reported (Fig. 5C). Quantification showed 54% degradation of ppGpp fifteen minutes after reversal of amino acid starvation (Fig. 5E, Table S3) while it was virtually undetectable after 1 minute in the *spoT* ^+^ strain (Fig. 5B, Table S2). When the pppGpp pool was elevated in the *spoT1* Δ*gppA* ::FRT/pRC*spoT* strain by SpoT depletion, the (p)ppGpp degradation was more severely impaired (Fig. 5D). There was only 22% degradation of ppGpp and 19% degradation of pppGpp after 15 minutes (Fig. 5E and Table S4). It is possible, the increase in (p)ppGpp pool from the turnover defect contributes to the allosteric activation of RelA and the growth arrest of *spoT1* Δ*gppA* strain.

### Synergistic inhibition of growth by ppGpp and pppGpp

As described above, the RelA-dependent growth arrest in the *spoT1*, *relA496*Δ::FRT Δ*spoT* and Δ*rlmD*::FRT Δ*spoT* strains were contingent on the loss of GppA function. Based on the principal biochemical activity of GppA, which is the conversion of pppGpp to ppGpp, and the elevated pppGpp pool observed during the growth inhibition, it was deduced as the primary determinant driving RelA activation, although it was less abundant than ppGpp during the growth arrest (Fig. 3B, lane 2; Fig. 4B, lane 13; Fig. 4D, lanes 2 & 4; Fig. 5D, lane 1). We asked, if elevated ppGpp, in the absence of pppGpp, inhibited growth, and if so, whether it was different from that seen in the additional presence of pppGpp. We took advantage of the phenotype of the Δ*rlmD*::FRT Δ*spoT* strain, which accumulated ppGpp but no detectable pppGpp during amino acid starvation (Fig. 2C (iii), lanes 6 to 8) and the ppGpp degradation rate was greatly diminished after the reversal of amino acid starvation (Fig. 6A), due to the absence of SpoT function. The ppGpp pool size was virtually unaltered for up to 60 minutes and reduced 28% and 37% respectively 120 and 180 minutes after the reversal of starvation (Fig. 6B and table S5). These results show SpoT as the primary ppGpp hydrolase.

**Figure 6.**
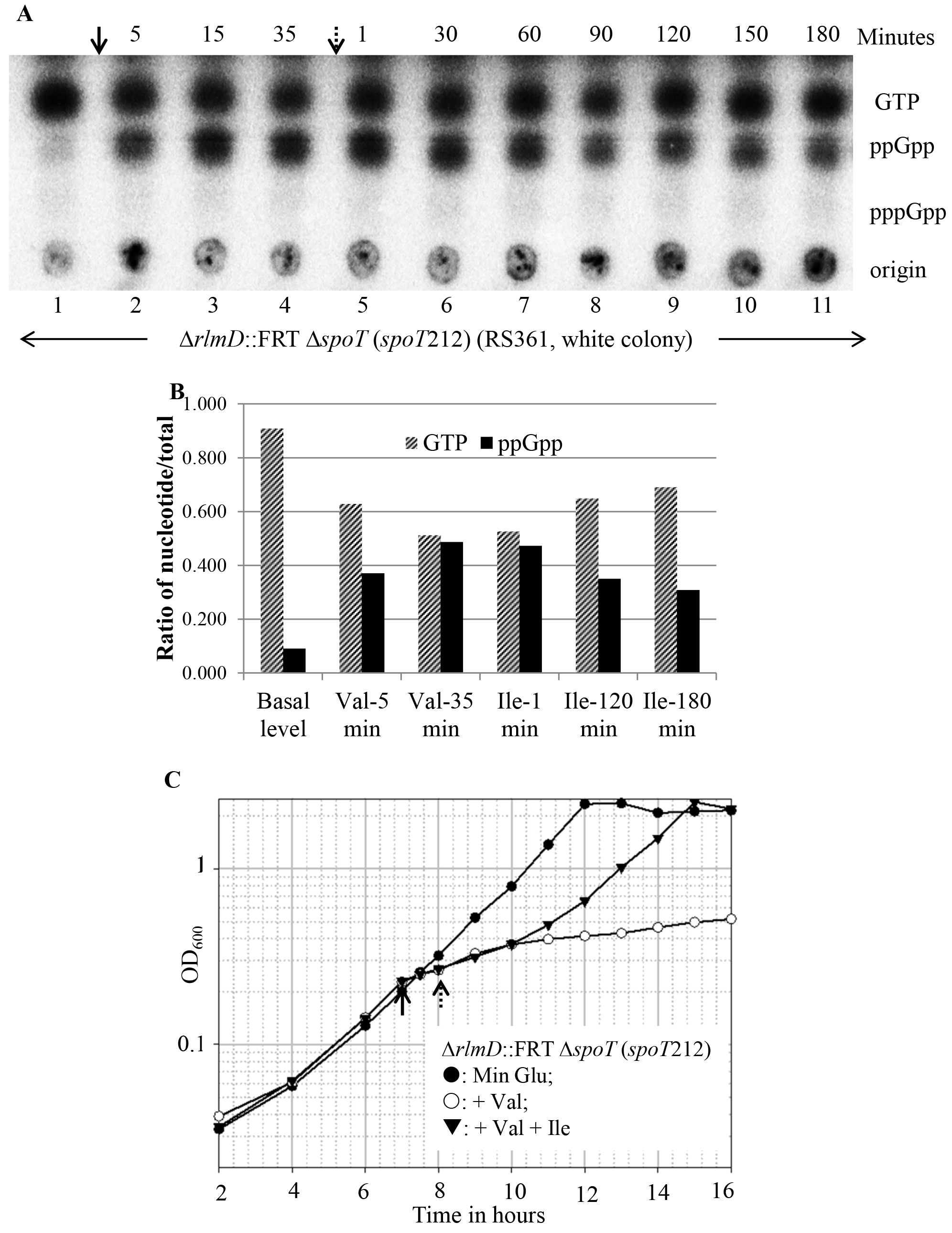
Accumulation of ppGpp is associated with transient growth arrest. **A.** TLC was performed for the strain Δ*rlmD* Δ*spoT* (RS361, white colony); growth and sampling are as described for Fig. 5A. **B.** The nucleotide pools were quantified from the TLC shown in Fig. 6A as described in Fig. 5B. **C.** Growth inhibition after valine addition (arrow, solid line) and recovery after isoleucine addition (arrow, dotted line) were determined for the Δ*rlmD* Δ*spoT* strain (RS361, white colony).

Growth arrest and recovery associated with amino acid starvation and its reversal were monitored in the Δ*rlmD*::FRT Δ*spoT* mutant and compared with that seen in the wild-type, *spoT1*, and Δ*rlmD*::FRT strains. As expected, growth ceased and resumed immediately following amino acid starvation and its reversal in the wild-type strain (Fig. S8A). In the *spoT1* strain, the starvation-induced growth arrest was similar to the wild-type, but growth resumed after a lag of ~30 minutes following the reversal of starvation (Fig. S8B), which correlated with the reduced rate of ppGpp degradation observed in the *spoT1* mutant (Fig. 5C). Isoleucine starvation in the Δ*rlmD*::FRT Δ*spoT* strain caused immediate growth arrest, but after the reversal of starvation, growth inhibition persisted for 120 minutes and then the growth resumed. After 120 minutes, the ppGpp pool was ~54% of GTP (Fig. 6B, 6C & Table S5) and this can be taken as the lowest pool size at which ppGpp conferred growth inhibition. One hour after growth resumed, that is, 180 after isoleucine addition, the ppGpp pool was ~45% of GTP (Fig. 6C and Table S5). Isoleucine starvation and its reversal in the Δ*rlmD*::FRT strain, which did not accumulate ppGpp (Fig. 2C(iii), lanes 2 to 4), produced growth arrest and reversal similar to the wild-type strain (Fig. S8C), and indicated that the accumulation of ppGpp was responsible for the transient growth arrest in the Δ*rlmD*::FRT Δ*spoT* strain. When growth inhibition from ppGpp and (p)ppGpp accumulation was compared, it was more severe when both pppGpp and ppGpp are present. For instance, in the *spoT1* Δ*gppA*/pRC*spoT* strain, where the growth arrest was severe and prolonged following SpoT depletion, ppGpp and pppGpp accumulated to 37% and 13% of GTP respectively after SpoT depletion (Fig. 5D, lane 1, table S4). These results showed that the growth inhibition conferred by ppGpp in isolation was transient, and this was strongly accentuated in the presence of pppGpp. While we did not measure ppGpp levels beyond 180 minutes, the continued growth of the Δ*rlmD*::FRT Δ*spoT* strain suggested that there could progressive decrease in the ppGpp pool and that eventually it may be degraded, although very inefficiently, by hydrolases other than SpoT.

## Discussion

### Autocatalytic activation of RelA by pppGpp

This study was initiated to test the prediction that the stringent nucleotides would accumulate in the cell following the depletion of SpoT activity. While an increase in the ppGpp pool was observed, accumulation of pppGpp was not seen and furthermore the pppGpp levels was not completely restored by inactivation of the pppGpp hydrolase, GppA. The results indicated that the degradation of pppGpp was enhanced during SpoT depletion and this was necessary to prevent RelA activation. RelA activation by pppGpp was clearly manifest in the *relA496*Δ Δ*spoT* and Δ*rlmD*::FRT Δ*spoT* strains, where, the slightly elevated ppGpp pool (Fig. 2C(i) lane 5 and 2C(iii) lane 5), increased several-fold after inactivation of GppA and pppGpp synthesis was evident (Fig. 4D lane 2 and 4). Since the (p)ppGpp pool increased in the absence of SpoT protein, and therefore totally independent of the hydrolase or synthase functions of SpoT, the increase in (p)ppGpp pool can be attributed entirely to the activation of RelA by the elevated basal pppGpp pool (from the loss of GppA activity). While the increase in the pppGpp pool may be important to initiate the allosteric activation of RelA, once initiated, the autocatalytic reaction may be sustained by both stringent nucleotides. ppGpp was reported to allosterically activate its own ribosome-dependent synthesis *in vitro* (Shyp *et al.*, 2012). However, given that a C-terminal His_6_-tagged protein was used in that study and RelA’s C-terminus resides well inside the ribosomal complex, it was likely that the functionality of the protein was compromised. In a recent report using untagged RelA protein, it was shown that pppGpp as an allosteric regulator and GDP as substrate synergize for the maximum enzymatic activity of RelA *in vitro* (Kudrin *et al.*, 2018). Allosteric activation of Rel protein by pppGpp was not restricted to full-length RelA, it has also been noted for RelQ, an RNA binding small alarmone synthase from *Enterococcus faecalis* (Beljantseva *et al.*, 2017) and as well as the small alarmone synthase, SAS1, from *Bacillus subtilis* (Steinchen *et al.*, 2015). While RelA is catalytically active as a monomer, SAS1 functions as a homotetramer and tetramerization was found to be important for the allosteric activation by pppGpp. A pppGpp binding site at the C-terminal region of the Rel enzyme from *Mycobacterium smegmatis* has been reported (Syal *et al.*, 2015). The physiological relevance of these findings remain unclear.

Our results provide experimental evidence, the first to our knowledge, that at least two levels of positive control on RelA activation exist *in vivo* (Fig. 7). In addition to the well-studied ribosome-dependent activation of the RelA catalytic activity by the hungry codons, there is a positive regulation of the RelA activity by pppGpp and which is likely to be an allosteric activation. While the activation from amino acid starvation requires RelA interaction with the ribosome the latter was also evident in the absence of ribosome binding (Fig. 4D). Further studies are needed to understand the physiological significance of the pppGpp mediated allosteric activation of RelA. By its very nature, an autocatalytic reaction would result in the generation of a bistable switch (Dubnau and Losick, 2006). We propose fluctuations in the SpoT and GppA hydrolase activities could contribute to the non-uniform (p)ppGpp synthesis in the cells within a population. It has been noted that the formation of persister cells within a population is associated with these cells having very high levels of (p)ppGpp (Harms *et al.*, 2016).

**Figure 7.**
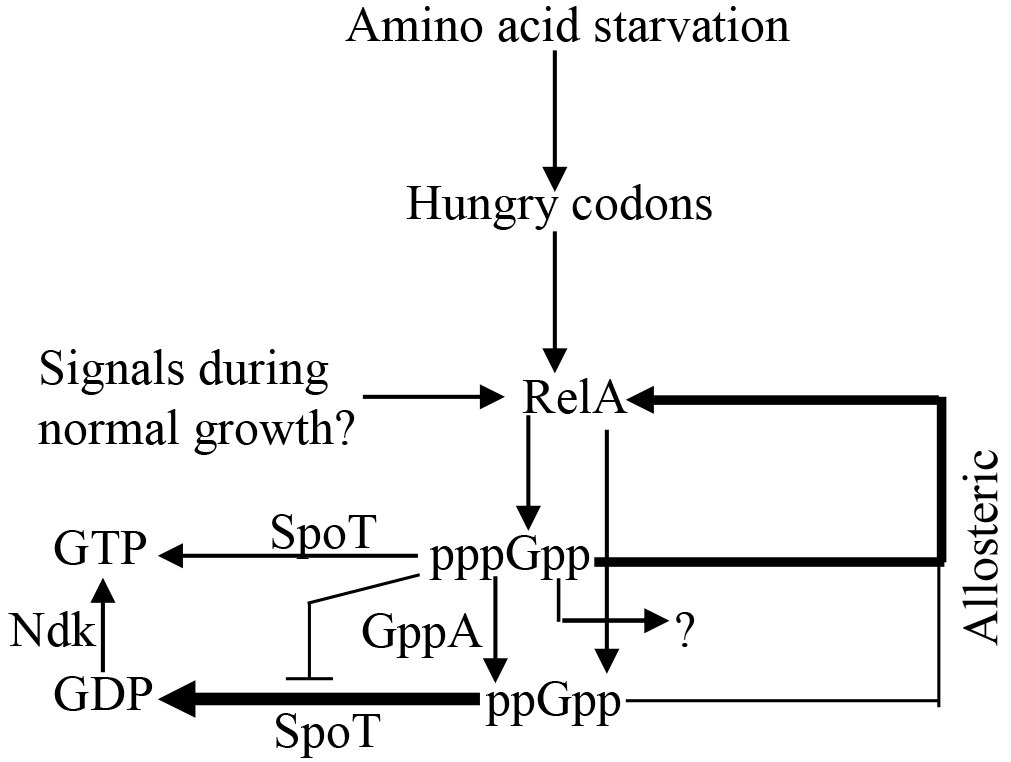
A model based on the results of this study. RelA activation from amino acid starvation is well established while the signals responsible for basal RelA activity are not clear. The stringent nucleotides generated are turned over by the SpoT and GppA hydrolases. When the SpoT hydrolase activity is lowered, two or more activities prevent the accumulation of pppGpp, one is GppA, the nature of the other one is unclear. The GppA and SpoT hydrolase activities together prevent the activation of RelA-mediated stringent response and growth arrest. The preeminent contribution of pppGpp to the activation of stringent response is because of the inhibition of the SpoT mediated degradation of (p)ppGpp and the allosteric activation of RelA (thick arrow); ppGpp may also allosterically activate RelA. See text for details. Ndk - Nucleoside diphosphate kinase. An arrow indicates activation or products of a biochemical activity, a line ending with a ‘┴’ indicates inhibition. A thick arrow refers to the strength /importance of the pathway.

### SpoT and GppA hydrolases - the negative regulators of the stringent response

Genetic evidence was used to postulate that the essential function of SpoT was the degradation of (p)ppGpp synthesized by RelA (Xiao *et al.*, 1991) in response to signal(s) the nature of which is not well understood. To our knowledge, this was not experimentally tested. Our results are in accord with this postulation but have revealed additional layers of regulation. While (p)ppGpp accumulation was anticipated from the depletion of SpoT, only ppGpp accumulation was observed, a phenotype reminiscent of that seen following amino acid starvation in the *spoT1* mutant. Although the molecular basis for this response is not clear, our results have eliminated the possibility that the pppGpp depletion arose solely from the activation of GppA, because, if that were to be the case, the pppGpp pool size would be completely restored in the Δ*gppA* background. However, the pppGpp pool observed in the Δ*gppA spoT1* and Δ*gppA* Δ*spoT* strains were less than that observed in the Δ*gppA* mutant (Fig. 3B lane 2, Fig. 4B lanes 10 - 16). Either the decreased synthesis of pppGpp or an increased degradation of pppGpp by one or more pppGppase, other than GppA, that was reported (Somerville and Ahmed, 1979) and references therein) may contribute to the depletion of pppGpp pool in the *spoT* hydrolase mutant. We favor the latter possibility, because, increasing the GTP pool did not increase the pppGpp synthesis (Fig. S6). Although the molecular mechanism is unclear, decrease in the pppGpp pool was an important adaptation needed to support the growth of strains deficient in *spoT* hydrolase activity.

Negative regulation of the stringent response by SpoT, consequent to amino acid starvation was evident in the Δ*rlmD*::FRT background (a hypomorphic allele with reduced expression of full-length RelA protein). Apparently, when there are fewer RelA molecules, signaling through the hungry codons was insufficient to amplify the (p)ppGpp pool faster than their turn over by the SpoT hydrolase activity (Fig. 2C (iii)). Further studies are needed to establish and understand the physiological relevance of such a regulation in the *relA*^+^ background. These results do not support the observations made *in vitro* that the uncharged tRNA’s inhibit the SpoT hydrolase activity (Richter, 1980), perhaps because the interaction between SpoT and uncharged tRNA possible *in vitro* may not occur in the cellular milieu. We propose, this kind of regulation could ensure that the amplification of the stringent response happens after a quorum of hungry codons is available to be sensed by RelA and tight negative regulation is maintained below the quorum through the SpoT and GppA hydrolase activities that lower the basal (p)ppGpp pool and prevent the activation of RelA by the nucleotides. Since, (p)ppGpp accumulation signals stress, such a regulation would ensure stress related adaptations are not provoked in response to minor fluctuations in the amino acid pool but only when there was sufficient starvation to allow a quorum of hungry codons to accumulate. When the rate of RelA-mediated (p)ppGpp synthesis exceeded the rate of SpoT/GppA mediated hydrolysis, the (p)ppGpp concentration can increase rapidly aided by the allosteric activation of RelA by pppGpp.

### Effect of (p)ppGpp versus ppGpp on growth

(p)ppGpp synthesis leads to growth inhibition. It has been reported that, in the absence of stress, progressively increasing the basal ppGpp level using hydrolase deficient *spoT* mutants progressively increased the growth inhibition (Fiil *et al.*, 1977; Sarubbi *et al.*, 1988; Xiao *et al.*, 1991). As noted previously (Laffler and Gallant, 1974; Fiil *et al.*, 1977) and as well as in this study, reduction in the *spoT* hydrolase activity decreased the pppGpp pool and therefore, the slow growth of the *spoT* hydrolase mutants are primarily due to elevated ppGpp pool and despite the reduced pppGpp pool. Since the elimination of the GppA activity increased the pppGpp pool in the SpoT hydrolase deficient strains (Fig 3B, 4B, and 4D), we compared the relative growth inhibition conferred by ppGpp with that of (p)ppGpp. While (p)ppGpp accumulation conferred severe and prolonged growth inhibition (Figs. 4A, 5D lane 1, Table S4) at comparable concentration, growth inhibition by ppGpp was transient (Figs. 6A lanes 9 - 11, 6B, 6C and Table S5). We propose, this could be because the two nucleotides regulate cellular functions with different efficiencies and which then synergistically contribute to growth inhibition. For example, it was reported that the inhibition of replication elongation was more pronounced when the basal pppGpp pool was increased (Denapoli *et al.*, 2013) while RNAP activity was more strongly inhibited by ppGpp (Mechold *et al.*, 2013). Our results also show that an important function of GppA is to lower the pppGpp pool and alleviate the gratuitous activation of RelA especially under conditions that lower the SpoT hydrolase activity. In other words, the SpoT and GppA hydrolases together maintain a low pppGpp pool so as to prevent the accumulation of (p)ppGpp through the autocatalytic amplification of basal RelA activity. The molecular mechanism of allosteric activation of RelA by pppGpp is being investigated.

## Experimental Procedures

### Growth conditions

LB medium (1% tryptone, 0.5% yeast extract, 1% NaCl), MOPS buffered minimal medium (Neidhardt *et al.*, 1974) or minimal A medium (Miller, 1992) were used, the latter was supplemented with 0.5% glucose and 20 amino acids, each at a final concentration of 40 μg ml^−1^. In plates, glucose and cas-amino acids were supplemented at 0.2% final concentration. Antibiotics and the concentrations at which they were used are, ampicillin (Amp), 50 μg/ml, kanamycin (Kan), 25 μg ml^−1^; tetracycline (Tet), 10 μg ml^−1^ and chloramphenicol (Cm), 15 μg ml^−1^. Isopropyl β-D-thiogalactopyranoside (IPTG) and 5-Bromo-4-chloro 3-indolyl-β-D-thiogalactoside (X-gal) were used at a final concentration of 1 mM and 50 μg ml^−1^, respectively.

### Construction of strains and plasmids

The strains used in this study are derivatives of the *E. coli* K12 strain MG1655, referred as wild-type. Strains and primers are listed in Table S1. Mutations were introduced by P1vir mediated transduction using standard protocols. Gene deletions have been sourced from the Keio collection (Baba *et al.*, 2006) and when required, the kanamycin resistance cassette was flipped out using FLP recombinase expressed from a pCP20 plasmid (Cherepanov and Wackernagel, 1995). The plasmid pRC*spoT* is from the lab collection (Nazir and Harinarayanan, 2016) and was constructed from pRC7 (Bernhardt and De Boer, 2004). Deletions in the *relA* gene on the chromosome, to generate *relA*455Δ::Kan and *relA*496Δ::Kan, wherein all codons after 455 and 496 were deleted and replaced with a TAG stop codon and the kanamycin cassette from pKD13 (Datsenko and Wanner, 2000) was performed by recombineering (Thomason *et al.*, 2014). Forward primers JGOrelA496aaPS4, JGOrelA455aaPS4, and the reverse primer JGOrelAPS1 were used. These constructs generated were verified by sequencing. The transposon insertions *relA*::Tn*10*dTet and *rlmD*::Tn*10*dKan were mapped by inverse PCR (Higashitani *et al.*, 1994). The β-galactosidase reporter fusions *relA’-lac* and *rlmD’-lac* within the *relA* and *rlmD* genes respectively, were constructed using the knockout alleles available in the Keio collection and plasmid pKG137 using a published protocol (Ellermeier *et al.*, 2002). The new junctions generated were verified by sequencing.

### Plasmid loss and viability measurements

Using a published assay (Bernhardt and De Boer, 2004) the ability of strains to grow following loss of the single-copy, unstable, amp^r^-plasmid pRC*spoT* was assessed in the *Δlac* genetic background. The plasmid carries the *lacZ* gene. Strains containing the plasmid pRC*spoT* were grown overnight in LB broth containing ampicillin and IPTG, the latter used at a final concentration of 1 mM unless mentioned otherwise. Overnight cultures were washed with minimal A medium and serially diluted to 10^−6^ dilution. Dilutions were spread on IPTG X-gal containing plates to obtain ~300 colonies and incubated for 24 to 72 hours at 37°C depending on the growth medium. Blue and white colonies indicating cells that retained and lost the plasmid respectively were counted and expressed as the percentage of white colonies. White colonies were tested for growth on the same medium to confirm viability.

### The efficiency of plating determination

Strains were grown overnight in the permissive medium. Overnight cultures were washed with minimal A and serially diluted to 10^−6^ dilution. From each dilution, 10μl was spotted, allowed to dry and plates incubated at 37°C. To test for RelA function or elevated (p)ppGpp level, strains were spotted on minimal A medium containing glucose with or without the amino acids serine (S), methionine (M) and glycine (G) (100 μg/ml each). The growth of the *relA* mutant was inhibited in the presence of SMG (Uzan and Danchin, 1976).

### Depletion of SpoT

In strains carrying the single-copy plasmid pRC*spoT*, the chromosomal *spoT* gene was replaced with the Δ*spoT*::Cm or the Δ*spoT* allele. SpoT expression in pRC*spoT* was induced using IPTG as it was expressed from the *lac* promoter and the plasmid also carried the *lacI* gene. Cultures grown overnight in the presence of ampicillin and IPTG (1mM) in LB or MOPS buffered medium containing glucose and 20 amino acids, were washed and subcultured at different dilutions in appropriate liquid medium or plates containing ampicillin and different concentrations of IPTG or without IPTG.

### (p)ppGpp estimation by thin layer chromatography

(p)ppGpp estimations were carried out by growing cultures in MOPS buffered medium containing 0.5% glucose and when necessary the 20 amino acids were added, each to a final concentration of 40 μg/ml. Overnight cultures and the initial growth following dilution were carried out in the presence of 1.32 mM K_2_HPO_4_. At an A_600_ of ~ 0.4 to 0.5, the cultures were diluted 10-fold into the low phosphate medium (0.4 mM K_2_HPO_4_) and allowed to undergo at least two doublings in the presence of 100-200 μCi/ml of ^32^P-H_3_PO_4_ before sample collection began at ~ 0.2 A_600_. An unlabelled culture was used to monitor A_600_ at periodic time intervals. Valine was added at 100 μg/ml to induce isoleucine starvation and which was reversed by adding isoleucine at 100 μg/ml. Samples were collected in tubes containing an equal volume of 2N HCOOH kept chilled on ice. The samples were subjected to three cycles of freeze-thaw and centrifuged at 10000 rpm for 5 minutes at 4°C and 10 μl of the supernatant was applied on PEI cellulose sheets and resolved in 1.5M KH_2_PO_4_, pH3.4. The nucleotide spots were visualized by phosphorimager (Typhoon FLA 9500) and quantified using multi-gauge software (Fujifilm). The values are expressed as the ratio of a nucleotide (GTP or ppGpp or pppGpp) to total (GTP + ppGpp + pppGpp).

### Growth kinetics of amino acid starvation and recovery

Overnight cultures were grown in minimal A glucose medium and sub-cultured in the same medium in three separate flasks. The cultures were incubated at 37 °C, 200 rpm and growth was monitored periodically. At OD_600_ between 0.15 - 0.25, valine was added to two of the cultures and 1 hour later isoleucine was added to one of the cultures having valine. Growth kinetics in the three flasks was compared.

## Acknowledgment

We acknowledge strains used from the Keio collection (NBRP, Japan). We thank members of the Laboratory of Bacterial Genetics for their suggestions. We thank Mike Cashel for providing strains used in this study and suggestions. This work was primarily funded by the Centre of Excellence in Microbial Biology - Phase 2 research grant of the Department of Biotechnology, Government of India. R.S is a recipient of DBT-JRF fellowship.

## Supporting information

**S1 Table.** List of strains, plasmids, and primers.

**S2 Table.** The ratio of the nucleotide to total in the WT and Δ*gppA* mutant following amino acid starvation and reversal of starvation.

**S3 Table.** The ratio of the nucleotide to total in the *spoT1* mutant following amino acid starvation and reversal of starvation.

**S4 Table.** The ratio of the nucleotide to total in the *spoT1* Δ*gppA*/pRC*spoT* strain after SpoT depletion and followed by amino acid starvation and reversal of starvation.

**S5 Table.** The ratio of nucleotides to total in the Δ*rlmD*::FRT Δ*spoT* strain following amino acid starvation and the reversal of starvation.

## SI references

**Fig. S1.** SpoT depletion in the Δ*relA* Δ*spoT*/pRC*spoT* strain. The Δ*relA* Δ*spoT* (*spoT212*)/pRC*spoT* (AN120) strain cultured overnight in MOPS medium containing glucose, 20 amino acids, ampicillin and 1mM IPTG was washed and sub-cultured at the dilution indicated in the same medium (•) or without IPTG (O).

**Fig. S2.** RelA-dependent accumulation of ppGpp but not pppGpp in the *spoT1* strain. *spoT1* (RS24) and Δ*relA*::Kan *spoT1* (RS485, white colony) strains were cultured in MOPS glucose medium and isoleucine starvation was induced with valine.

**Fig. S3.** SMG resistance of the hypomorphic RelA alleles is modulated by SpoT function. Saturated cultures were washed, serially diluted and spotted on plates with or without SMG (SMG refers to serine, methionine, and glycine; see methods) and photographed after 20 hrs at 37^°^C. Strains and their relevant genotypes are indicated. * growth in the presence of SMG was retarded but not abolished.

**Fig. S4.** The effect of *spoT* and *gppA* deletions on the growth of the *relA1* strain. Growth was monitored by streaking the indicated strains on Minimal glucose SMG plates and incubating them for the time indicated (A) or in other plates as indicated (B). LB and Minimal Glucose CAA plates were incubated for 24 hours and Minimal glucose plates were incubated for 48 hours. The strains were MG1655, *relA1* (RS39), *relA1* Δ*spoT*(RS31 white colony), and *relA1* Δ*spoT* Δ*gppA* ::Kan (RS35 white colony).

**Fig. S5.** Expression levels of *relA ‘-lac* and *rlmD’-lac* fusions in LB. β-galactosidase assay was carried out from mid-log phase cultures of the strains RS692 (*relA-lac*) and RS693 (*rlmD’-lac*) grown in LB. The β-galactosidase activity in Miller units was plotted on a log scale. Values obtained from three independent experiments were used to calculate the mean and standard deviation.

**Fig. S6.** The absence of pppGpp accumulation in the *spoT1* strain is not corrected by an increase in the GTP level. The *spoT1 gsk3* (RS697 white colony) strain was cultured in MOPS glucose medium containing histidine, tryptophan (each at 100 μg/ml) without guanosine (lanes 1 to 4) and with guanosine (lanes 5 to 8). Isoleucine starvation was induced by the addition of valine (indicated by arrow). Samples were collected immediately before valine addition or subsequently at the time indicated.

**Fig. S7.** SpoT depletion in spoT207::Cm Δ*gppA* ::Kan/pRC*spoT* (RS206). The spoT207::Cm Δ*gppA* ::Kan/pRC*spoT* (RS206) grown in MOPS medium containing glucose, 20 amino acids, ampicillin and 1mM IPTG was washed and sub-cultured in the same medium (•) or without(O) IPTG at the dilution indicated. Growth after the sub-culture into the medium containing ^32^P-H_3_PO_4_ is shown. The arrows refer to the points at which samples were collected for the TLC shown in Fig. 3B.

**Fig. S8.** Growth inhibition by valine and the recovery after isoleucine addition were assayed in **A.** MG1655, **B.** *spoT1* (RS24), and **C.** Δ*rlmD* (RS316). Valine (solid arrow) and isoleucine (dashed arrow) were added as indicated.

